# Rapid detection, quantification and speciation of *Leishmania* using real-time PCR and DNA sequencing at the rRNA Internal Transcribed Spacer 2

**DOI:** 10.1101/2024.05.20.595045

**Authors:** Andrea Paun, Michael E. Grigg

## Abstract

The ability to discriminate infection between closely related *Leishmania* species within the *Viannia* species complex, specifically *L. braziliensis, L. guyanensis* and *L. panamensis* is critical to inform the clinical diagnosis and determine the most efficacious treatment modality. We designed a nested primer set targeting the rRNA Internal Transcribed Spacer 2 (ITS2), located on Chromosome 27, to distinguish among all human infective *Leishmania* species. Separate nested and single primer pairs were developed for conventional and quantitative PCR approaches respectively. Species-specific single nucleotide polymorphisms and indels located within the PCR products were identified by Sanger sequencing. This single locus approach provides a sensitive and specific platform to identify the species of *Leishmania* causing infection.

## Introduction

Leishmaniasis is a spectrum of diseases resulting from infection with parasites within the genus *Leishmania* following the bite of an infected insect vector (1-6). Leishmaniasis is caused by more than 20 species of *Leishmania* and is endemic in tropical and sub-tropical regions where vector species are endogenous. Clinical presentation can range from both self-healing and persistent cutaneous lesions, mucocutaneous lesions to potentially fatal visceral infection. The parasite species, geography and the host’s genetics and immune status can all affect symptoms, making a species level diagnosis potentially complicated. Areas of endemicity often have several species of *Leishmania* present and as treatment options have species-dependent efficacy, the correct speciation plays an important role in determining treatment success (7-10). This is of particular relevance in travel-acquired infections, as patients can acquire infections with species that are different to their home country or return to countries where Leishmaniasis is not present and therefore is not commonly diagnosed by physicians. Rapid and accurate diagnosis and species identification can therefore facilitate clinical management, leading to improved patient outcomes.

Historically a number of methods have been used to diagnose Leishmaniasis including, most commonly, direct microscopic examination of clinical specimens and culture, neither of which allows for speciation. Isoenzyme analysis has long been the standard for speciation; however, this relies on the ability to grow a specimen in culture and is technically demanding and time consuming. More recently diagnosis has been facilitated by advances in molecular biology and the PCR amplification and sequencing of various genes has allowed for significantly faster speciation of *Leishmania* parasites directly from clinical samples. A range of genes have been utilized for speciation including *7SL, HSP70*, and *CytB*, amongst others (11-16).

The Internal transcribed spacer 2 (ITS2) has previously been identified as a region of utility for speciation and has been demonstrated to provide speciation for both Old World and New World *Leishmania* species (17, 18). Located on Chromosome 27, the region is flanked by highly conserved 5.8S and 28S ribosomal RNA subunit genes. Conserved primers selected within these two genes were generated to amplify across the more polymorphic and size variable ITS2 locus to produce a pan-genus typing strategy that resolves the species of *Leishmania* causing infection. Sequences derived from this approach were used to generate another conserved primer set that is anchored within the ITS2 locus to develop a real-time PCR assay that amplifies a 280-300 bp product to quantitate and speciate by Sanger sequencing the *Leishmania* species present in clinical samples. This approach allows for greater flexibility in both clinical and research settings, and can be combined with other sensitive primer sets that amplify genes located with the organellar kDNA genome (*i*.*e*., *7SL* gene) to detect with high sensitivity and specificity all major species of *Leishmania*.

## Materials and Methods

### Specimen collection, handling, and parasite culture

*Leishmania* parasites were grown in Complete M-199 media (Sacks and Melby, 2001). Reference strains, listed in Table 1 were obtained from American Type Culture Collection (Manassas, VA).

**Table 1.**
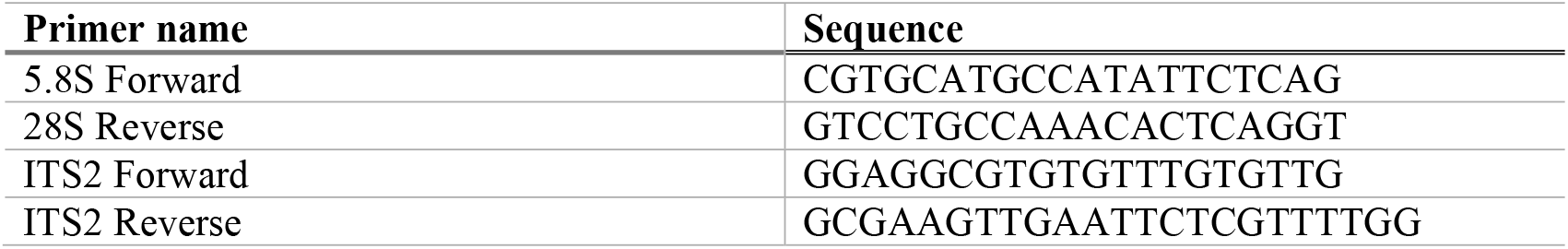
Primers.

**Table 2.**
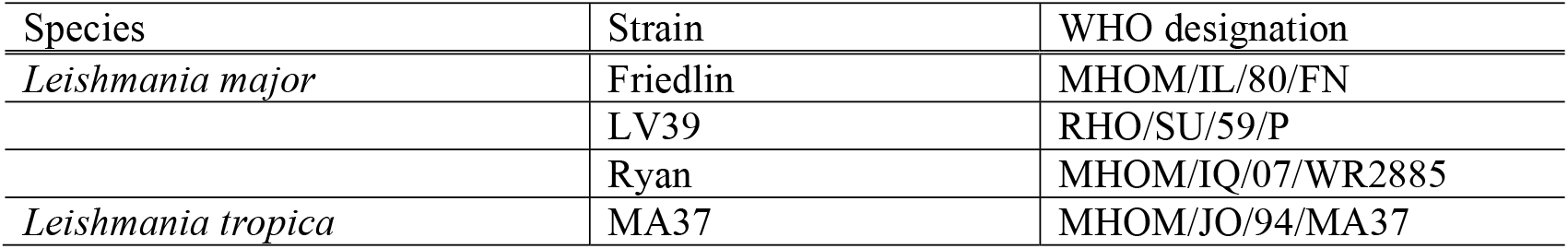
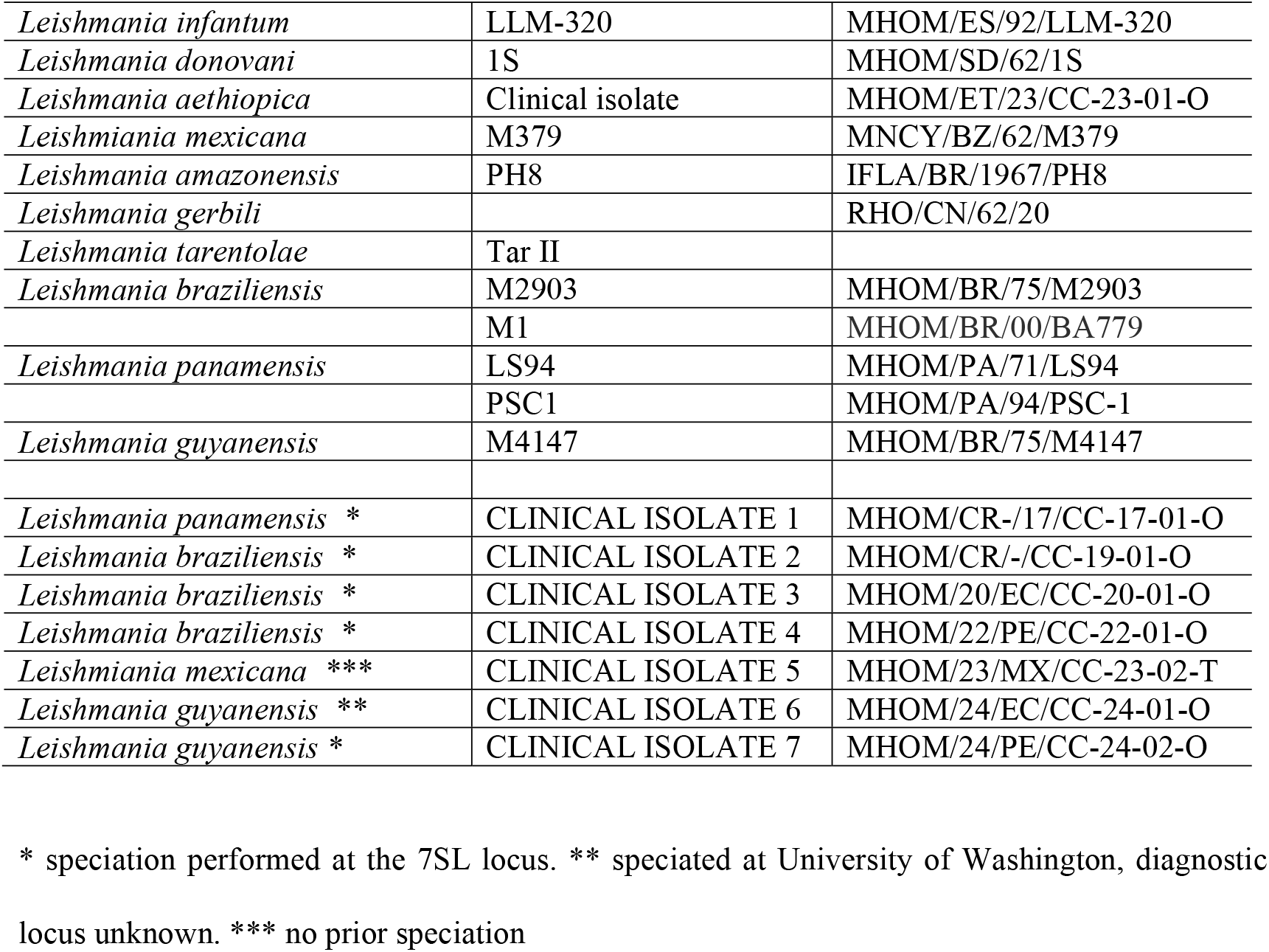
Reference and clinical isolates.

Clinical specimens were obtained from subjects enrolled in an Institutional Review Board-approved leishmaniasis protocol (No. 01-I-0238, study ID NCT00344188) at the National Institutes of Health Clinical Center (Bethesda, Maryland, USA). Subjects provided written informed consent for the collection and use of research specimens. Samples were kept in sterile saline and clinical isolate outgrows were established, as previously described (Paun, 2022).

### DNA extraction, PCR and Sanger sequencing

For reference and outgrow strains, genomic DNA was obtained from 5ml of stationary phase culture of parasites using the DNeasy Mini Kit (Qiagen, CA) following manufacturer’s instructions. For PCR, reactions were conducted in a total volume of 20 µL containing 1 µL of sample DNA, 0.2 µM; of FWD and REV primers (Table 1) and 10 µL of 2x SybrGreen PCR Mastermix (Applied Biosystems) in PCR molecular grade water. PCR was performed using the following cycling conditions: initial denaturation 95°C for 10 minutes; followed by 30 cycles of 95°C for 30 s, 60°C for 30 s, 72°C for 60 s and a final extension phase of 72°C for 5 min. The amplified products were resolved on 1-8-3% agarose gels, as indicated, and visualized by staining with SybrSafe (LifeTech).

For real time PCR, reactions were conducted in a total volume of 25 µL containing 1 µL of sample DNA, 0.1 µM; of ITS2 FWD and ITS2 REV primers (Table 1), 5 µL of 5x KAPA HiFi GC buffer, 0.3 mM dNTP mix, and 0.5 U of KAPA HiFi DNA polymerase (Roche). PCR was performed using the following cycling conditions: warming to 50°C for 2 minutes, initial denaturation 95°C for 5 minutes; followed by 40 cycles of 95°C for 15 s and 60°C for 60 s, with a melt curve analysis performed at the end of cycling.

Initial speciation of clinical samples was conducted by the NIH Department of Laboratory Medicine Microbiology Service using the kDNA *7SL* gene marker. DNA from punch biopsy specimens was obtained using the NucliSens kit and easyMAG (bioMerieux Inc., Durham, NC) following manufacturer’s instructions. Sequencing of the *7SL* gene was performed for speciation as described previously (Stevenson, 2010).

PCR products were purified using Ampure Beads (Beckman) following manufacturer instructions and Sanger sequenced at the RML Research Technologies Section, NIAID, NIH. Consensus sequences and alignments of the internal (qPCR) product were generated in Geneious Prime. MEGA-X was used to generate a phylogenetic tree using the Maximum Likelihood Method and Tamura-Nei model.

### Limit of detection (LOD) determination

The limit of detection was determined for the qPCR assay utilizing the internal primer set. This was done for both parasites/µL, counted by hemocytometer and ng/µL as determined by Qubit (ThermoFisher Scientific). Standard curves with a range of 10^6^-10^-1^ parasites/microliter and 10ng – 100fg/microliter were made by 10-fold serial dilution in PCR-grade water. No template controls (NTC) were included for both curves.

## Results

### Assay design and validation

The utility of the ITS2 region for speciation of *Leishmania* has been previously demonstrated using clinical samples in combination with PCR and Sanger Sequencing to resolve a limited number of species (17, 18).

In order to optimize the utility of this assay in both a research and clinical setting, we developed nested primers anchored in the 5.8S and 28S ribosomal genes that amplify across the more polymorphic and size variable ITS2 locus in order to detect all species of *Leishmania* of clinical significance. Nested primers were prioritized so that samples with a relatively low amount of parasite DNA present could undergo an additional round of nested amplification to increase the sensitivity of this assay to speciate the *Leishmania* parasite present (Fig. 1A). Specifically, external primers (5.8S FWD and 28S REV) were identified on Chromosome 27 within the highly conserved regions of the 5.8S and 28S (LSUα) ribosomal RNA genes respectively, allowing for amplification of the entire ITS2 sequence (Fig. 1A). The regions selected were highly conserved across all species of *Leishmania*, thus providing a pan-genus species-typing assay. These primers resulted in size variable amplification products of approximately 580-600bp for species in the *Viannia* complex, a larger fragment ranging from 750-800bp for Old World *Leishmania* and *Leishmania Mundinia*, and an approximately 900bp product for *Sauroleishmania* (Fig 1B). Sequence alignments identified multiple insertions and deletions that resulted in the diverse range of product sizes observed in all strains, except the much more conserved *Viannia* species.

**Figure 1.**
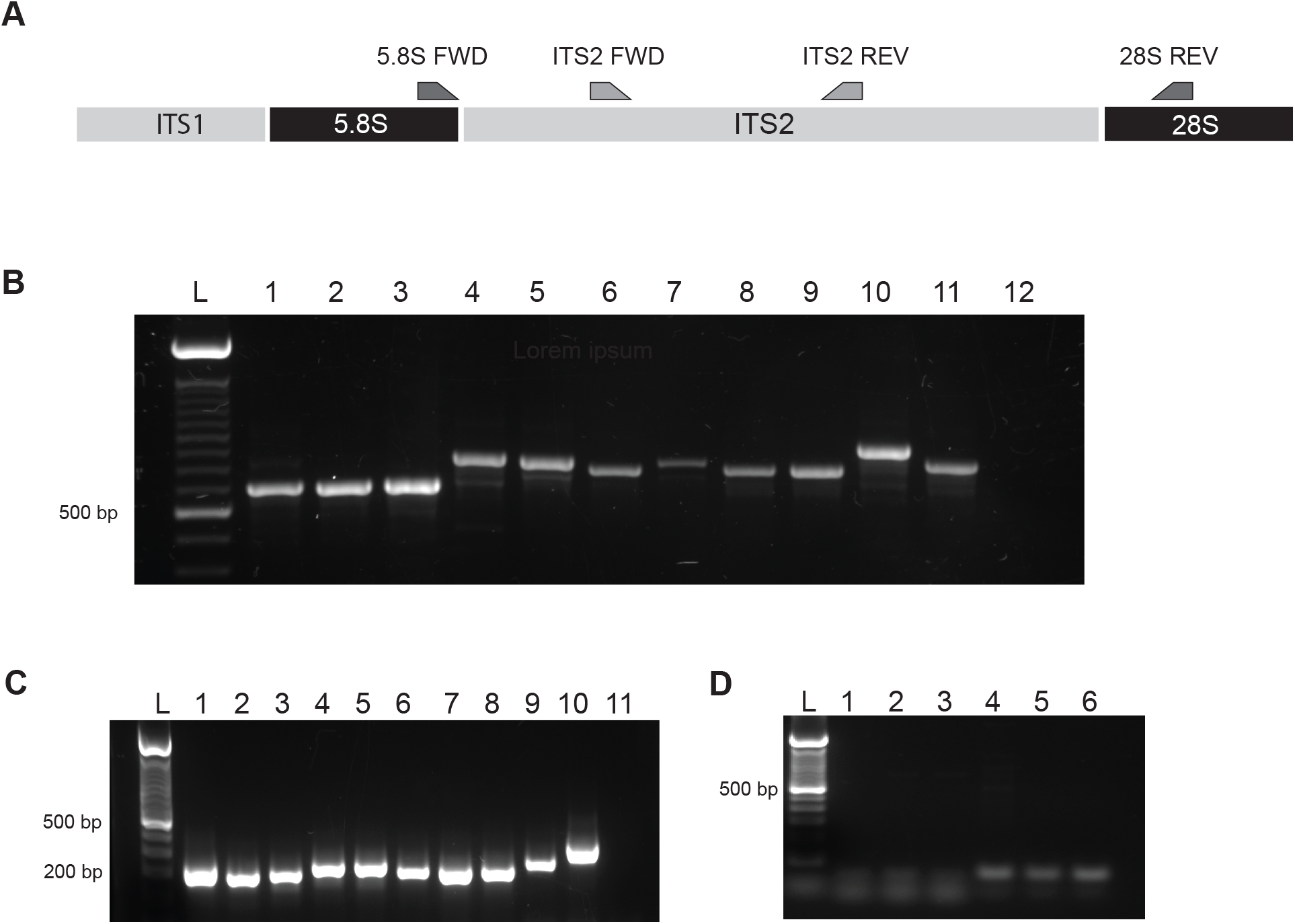
Molecular amplification of internal transcribed spacer 2. A. Schematic representation of the locus with the location of the primers. B. Agarose gel (1.8%) of the PCR products generated using the external primer set (5.8S FWD and 28S REV) for 1-*L. braziliensis M2903*, 2-*L. panamensis LS94*, 3-*L. guyanensis*, 4-*L. major Fn*, 5-*L. mexicana*, 6-*L. tropica*, 7-*L. donovani*, 8-*L. infantum*, 9-*L. amazonensis*, 10-*L. tarentolae*, 11-*L. gerbilli*, 12-NTC. C. Agarose gel (3%) of the internal primers set to be used for qPCR (ITS2 FWD and ITS2 REV) for 1-*L. braziliensis M2903*, 2-*L. panamensis LS94*, 3-*L. guyanensis*, 4-*L. major Fn*, 5-*L. mexicana*, 6-*L. tropica*, 7-*L. donovani*, 8-*L. infantum*, 9-*L. amazonensis*, 10-*L. gerbilli*, 11-NTC. D. Agarose gel (1.8%) of the PCR products generated using the external primer set for 1-*T. cruzi*, 2-*T. brucei*, 3-*T. gambiensi* and the internal primer set for 4-*T. cruzi*, 5-*T. brucei*, 6-*T. gambiensi*.

In settings where the quantitation of parasite DNA is also desirable, an internal primer set (ITS2 FWD and ITS2 REV) was identified in regions within ITS2 that showed conservation across all species tested. The amplicon generated was 280-335 base pairs in size (Fig. 1C) depending on the species present and possessed sufficient sequence variability to allow for the identification of all species included in the study (Supp. Fig 1). Importantly, this PCR product contains four diagnostic SNPs and four indels that allow for discrimination between *L. braziliensis, L. panamensis* and *L. guyanensis* by Sanger sequencing of the PCR product (highlighted, Supp. Fig 2). To determine species specificity, the primer sets were also tested for their ability to amplify the closely related species *Trypanosoma cruzi, T. brucei* and *T. gambiensi*. Neither primer pair was able to PCR amplify products from these species, indicating their exclusive specificity for parasites within the genus *Leishmania* (Fig. 1D).

**Figure 2.**
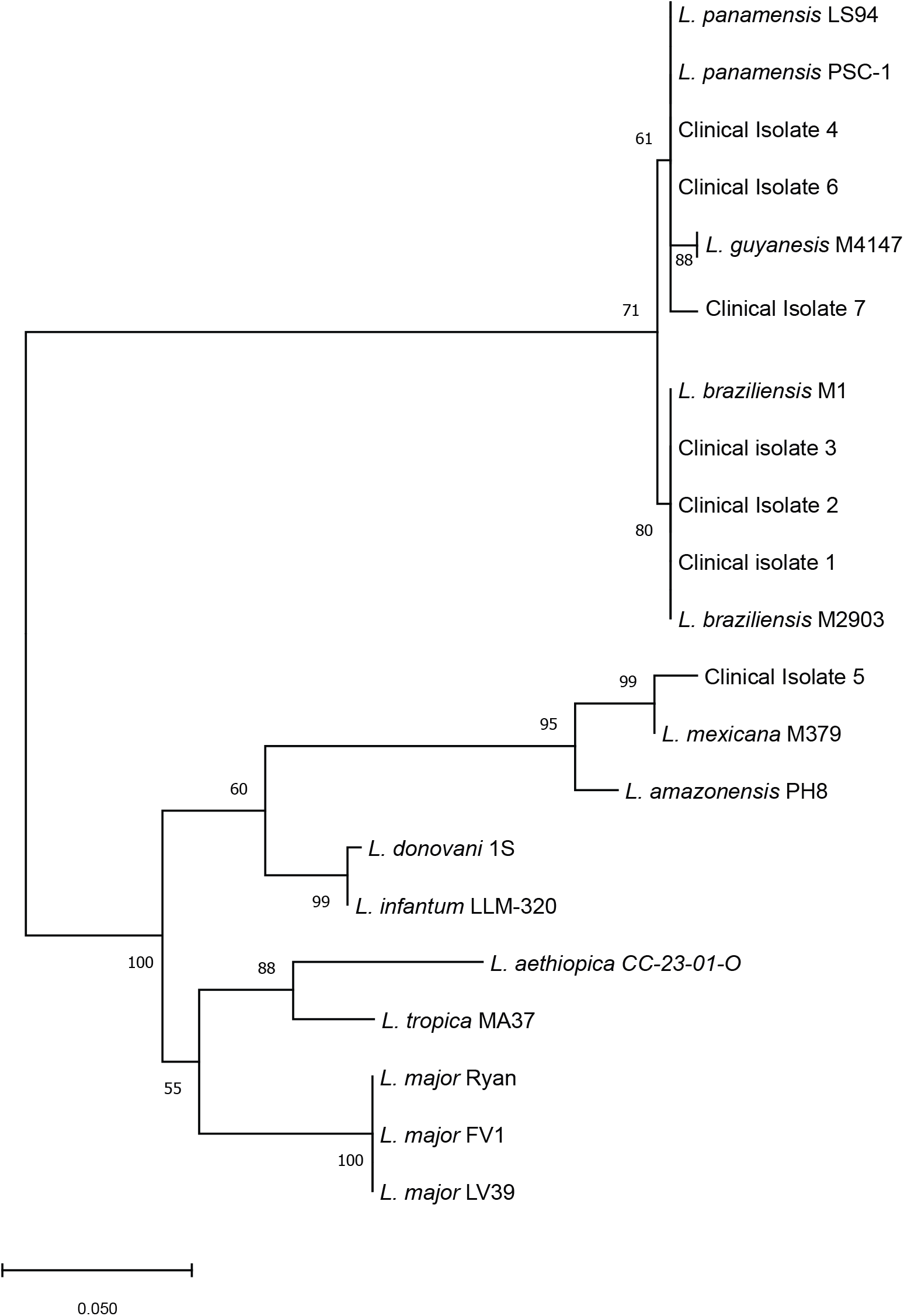
Phylogenetic tree. Phylogenetic tree of 16 reference isolates and 7 clinical isolates of *Leishmania spp*. was constructed by using the Maximum Likelihood Method and Tamura-Nei model in MEGA X with the sequences of the internal (qPCR) product of ITS2. The number on the branches represent the percentage of 1000 bootstrap samples supporting the branch. Only values >50 are shown.

### Limit of detection of qPCR assay

Clinical isolates received at NIH DLM are routinely analyzed at the 7SL gene, a marker in the organellar genome of *Leishmania*. To determine the sensitivity of the nuclear genome encoded ITS2 internal primer set, we compared the ITS2 assay against the 7SL assay. The limit of detection (LOD) of the qPCR assay was determined by titration of both quantitated parasites as well as the quantity of parasite DNA and compared directly to the 7SL gene (11, 12). For the purposes of comparison by qPCR, the primer set from Zelazny *et. al*. was utilized, it amplifies a 190bp product which is an optimal size for qPCR.

The ITS2 primer pair had an LOD equivalent to less than 1 parasite, which was more sensitive then the 7SL assay by approximately 1 log (Supp Fig. 3A). The addition of 30ng of human genomic DNA did not inhibit the sensitivity or specificity of the assay, as determined by C_T_ and melt curve analysis. Similarly, using a standard curve generated with defined quantities of parasite DNA, both the ITS2 and 7SL primer pairs were able to detect as little as 100fg/µl of *Leishmania* DNA, however, the C_T_ was approximately 4.2 C_T_ lower for ITS2 (Supp. Fig 3B). Therefore the ITS2 assay provides at least a log-fold greater sensitivity of detection when compared to the 7SL assay by both units of measure for quantification of parasite DNA.

### Speciation of clinical samples

The ability to speciate clinical specimens was examined using a set of seven clinical isolates provided by the NIH Clinical Center Department of Laboratory Medicine Microbiology Service, five had previously been speciated at the 7SL locus, two had insufficient parasite DNA present to allow for speciation by the 7SL assay (Clinical isolates 5 and 6), however Clinical Isolate 6 had previously been speciated at a different institution. Culture outgrows were obtained for all isolates with the exception of Clinical Isolate 5, because a tissue biopsy was not provided (19). DNA was extracted, the ITS2 FWD and REV primers were used to amplify a product that was Sanger sequenced. Sequence alignments (Supp Fig 2 – isolates 1-4, 6 & 7) and phylogenetic analysis were used to speciate the isolates (Fig 2). Clinical isolates 1-3 clustered with *L. braziliensis*, Clinical isolates 4 and 6 clustered with *L. panamensis*, and Clinical isolate 5 clustered with *L. Mexicana*. Clinical isolate 7 was found to cluster in a branch containing *L. panamensis* and *L. guyanensis* but was distinct from both species. In samples where both tissue DNA and culture-derived DNA were present, PCR-DNA Ssequencing yielded the same sequence for both samples, indicating that the presence of human genomic DNA or variable quantities of parasite DNA did not impair the specificity or sensitivity of the assay.

## Discussion

The ability to identify *Leishmania* spp. quickly and accurately as part of a clinical diagnosis is critical for the treatment plan and subsequent clinical outcome of patients. The clinical presentation of cutaneous lesions is very similar for many species, and with many endemic areas having more than one species of *Leishmania* the use of molecular diagnostics allows for rapid identification of the causative species. This is particularly important for *Leishmania Viannia* spp. where there is the potential for irreversible mucocutaneous tissue destruction. The utility of the ITS2 rRNA gene locus located on Chromosome 27 has previously been identified for its ability to differentiate among some species of *Leishmania* (17) as well as between New World and Old World species (18). These approaches used conventional PCR followed by Sanger sequencing on only a limited number of samples and species to compare with standard laboratory identification methodologies.

We sought to improve on the sensitivity and specificity of this locus to provide maximum utility in both clinical and research settings. Hence, we designed a nested set of pan-genus primers that amplified across the ITS2 region by using primers anchored in highly conserved regions of the flanking 5.8S and 28S rRNA subunit genes (Fig 1A). The difference in band size found between New and Old World species due to the presence of indels allowed for an immediate, albeit crude speciation after resolving DNA present by agarose gel electrophoresis. The internal primer set utilizes highly conserved regions found within the ITS2 rRNA locus that contains sufficient variation to accurately identify all tested species of clinical significance in a fragment of approximately 300 bp (Fig 1C, Supp Fig 1). Importantly we found no amplification of any of the *Trypanosoma* spp. tested for cross reactivity (Fig 1D) therefore providing a *Leishmania*-specific assay.

In some settings, it is desirable to use qPCR in order to determine the amount of parasite DNA present. This can be of particular importance as the parasite is not often uniformly distributed throughout the lesion and this information may be required to determine the suitability of a sample for further molecular study or for clinical reporting. The high copy number of the trans-ITS region (estimated at 20-40 copies, depending on species) therefore makes it an excellent target when high sensitivity is required (20). This is borne out by the LODs determined for the internal primer pair, with at least a log-fold greater sensitivity when compared against the 7SL locus (Supp Fig. 3), thus providing a greater sensitivity and likelihood of identifying the species present in the patient sample.

As there are strains of emerging clinical relevance within *Leishmania Mundinia*, such as *L. martiniquensis*, we included *L. gerbili* from the same species complex as a test for the ability of the primers to amplify within the *Mundinia* complex of strains. Although we were unable to obtain an isolate of *L. martiniquensis*, our analysis of deposited sequences in GenBank (CM030422.1) confirmed conservation of all primer binding regions and predicted a qPCR product of comparable size to that generated for *L. gerbili*, which clustered distinctly from *Leishmania* and *Leishmania Viannia* strains, suggesting the feasibility of using this region for future studies involving speciation of *Leishmania Mundinia* spp.

For species level identification of *Leishmania Viannia* spp., we confirmed the presence of the three SNPs identified by de Almeida et al., 2011, and that the unique *HphI* restriction site specific to *L. guyanensis* was present, and could conceivably be used to distinguish *L. guyanensis* infection from that of *L. braziliensis* or *L. panamensis*, as reported previously (17). We also identified an additional SNP and four indels that, when combined, gave a unique sequence profile for each of the *Leishmania Viannia* species within the internal ITS2 PCR product.

During assay development, the sensitivity and specificity of the nuclear genome ITS2 locus was compared against the mitochondrial (organellar) genome 7SL locus for diagnosis and speciation of infection. While the ITS2 locus was determined empirically to be more sensitive and specific than the 7SL locus, the utility of developing a two marker qPCR assay that combined both loci for clinical diagnosis was recognized. Inter-specific genetic hybridization among *Leishmania* species within the *Viannia* species complex is increasingly being reported, with mito-nuclear discordance a hallmark for detecting hybrids (21-28). Among the six clinical isolates speciated using the ITS2 locus herein, that were also sequenced at the 7SL locus, 4 isolates showed mito-nuclear incongruence. Furthermore, the allele identified at ITS2 for clinical isolate 7 appeared to be recombinant because it shared features with both *L. panamensis* (SNP pattern) and *L. guyanensis* (indel pattern). The allele at the mitochondrial 7SL marker was that of *L. guyanensis*, further supporting the tenet that this isolate was an interspecific recombinant, and would be an ideal candidate for whole genome sequencing. In aggregate, our data suggests that multilocus typing should be utilized in settings where this is feasible, rather than relying on a single locus to speciate *Leishmania Viannia* isolates.

In summary, we have developed a set of pan-genus *Leishmania*-specific nested PCR primers that allow for the speciation of all *Leishmania* spp. of current clinical relevance using a combination of PCR and Sanger sequencing. Further, the internal primer set is appropriate for settings where qPCR is used and contains sufficient sequence variation to speciate both Old World and New World species in both a sensitive and specific manner. This assay can be utilized in both research and diagnostic settings where the molecular characterization of parasites is performed.

## Acknowledgements

This research was supported by the Intramural Research Program of the National Institute of Allergy and Infectious Diseases. We thank Dr. Elise O’Connell and Dr. Janitzio Guzman of the Laboratory of Parasitic Diseases (LPD), NIAID, and the LPD clinical staff for their collection of patient samples. We thank Dr. Joshua Lacsina of the LMVR, NIAID for helpful discussion. Speciation was performed by the National Institutes of Health Department of Laboratory Medicine Microbiology Service.

## Figure legends

**Supplementary Figure 1**.

Alignment of the ITS2 PCR-amplified region from *Leishmania* spp. Representative sequence information from 1 isolate of each species is shown. Dots indicate identity with L. major sequence.

**Supplementary Figure 2**.

Alignment of the ITS2 PCR-amplified region from *Leishmania Viannia* spp. including clinical isolates. Representative sequence information from 1 isolate of each species is shown. Dots indicate identity with *L. braziliensis* M2903 sequence. Highlighted areas indicate diagnostic SNPs and indels.

**Supplementary Figure 3**.

Standard curves of ITS2 and 7SL shown as amplification plots and tables of Ct values for each concentration of *L. braziliensis* M2904 genomic DNA for A. Organisms per microliter or B. Quantity of genomic DNA per microliter.

**Supplementary Table 1**

Mito-nuclear incongruence between the 7SL and ITS2 genotyping loci for six clinical isolates. The following abbreviations were used to denote the species of the allele identified: Lbz, *Leishmania braziliensis*; Lp, *Leishmania panamensis*; Lg, *Leishmania guyanensis*. An asterisk (*) was used to identify a variant allele that possessed one or more private SNPs from the allele of the closest species.

